# Mapping of the *bs5* and *bs6* non-race-specific recessive resistances against bacterial spot of pepper

**DOI:** 10.1101/2022.09.26.509408

**Authors:** Anuj Sharma, Jian Li, Rebecca Wente, Gerald V. Minsavage, Upinder S. Gill, Arturo Ortega, C. Eduardo Vallejos, John P. Hart, Brian J. Staskawicz, Michael R. Mazourek, Robert E. Stall, Jeffrey B. Jones, Samuel F. Hutton

## Abstract

Bacterial spot caused by *Xanthomonas euvesicatoria* is a major disease of pepper (*Capsicum annuum* L.) in warm and humid production environments. Use of genetically resistant cultivars is an effective approach to manage bacterial spot. Two recessive resistance genes, *bs5* and *bs6*, confer non-race-specific resistance against bacterial spot. The objective of our study was to map these two loci in the pepper genome. We used a genotyping-by-sequencing approach to initially map the position of the two resistances. Segregant populations for *bs5* and *bs6* were developed by crossing susceptible Early CalWonder (ECW) with near-isogenic lines ECW50R (*bs5* introgression) or ECW60R (*bs6* introgression). Following fine-mapping, *bs5* was delimited to a ~535 Kbp interval on chromosome 3, and bs6 to a ~666 Kbp interval in chromosome 6 of pepper. We also identified 14 and 8 candidate resistance genes for *bs5* and *bs6*, respectively, based on predicted protein coding polymorphisms between ECW and the corresponding resistant parent. Mapping of *bs5* and *bs6* will facilitate their use in breeding programs through marker-assisted selection and is also a crucial step towards understanding the mechanisms of resistance.

**Key message:** Two recessive bacterial spot resistance genes were mapped in the pepper genome, which will facilitate their advancement in commercial pepper for management of all races of *Xanthomonas euvesicatoria*.

## Introduction

Pepper (*Capsicum annuum* L.) is an important solanaceous crop that is cultivated throughout the world. Bacterial spot of pepper (BSP) is a major disease responsible for loss of marketable yield in many pepper-growing regions (Osdaghi et al. 2021). The disease is manifested as dark brown necrotic lesions in all aerial parts of the plant. Foliar infection can lead to defoliation, which in turn leads to yield loss. The marketability of fresh fruits is also affected by the presence of scab-like symptoms or due to sun-scalding resulting from extensive defoliation (Ritchie 2000). The disease is caused by three species of *Xanthomonas* — *X. vesicatoria, X. euvesicatoria* (*Xe*), and *X. gardneri* (*Xg*) (Osdaghi et al. 2021). The management of BSP often relies on application of copper-based bactericides; however, the emergence of copper-tolerant strains has rendered this strategy unsustainable (Stall *et al*. 2009). Alternatively, host plant resistance has been applied as an effective, economical, and environmentally friendly way of managing BSP.

Most of the resistances deployed in modern agriculture involve a dominant resistance (R) gene which often belong to Nucleotide-Binding Leucine Rich Repeats (NLR) or Receptor-Like Kinase (RLK) protein family (Sharma et al. 2022a). Five dominant resistances have been reported against BSP — *Bs1* from *C. annuum* accession PI 163192 (Cook and Stall 1963), *Bs2* from *C. chacoense* PI 260435 (Cook and Guevara 1984), *Bs3* from *C. annuum* PI 271322 (Kim and Hartmann 1985), *Bs4C* from *C. pubescens* PI 235047 (Sahin and Miller 1998), and *Bs7* from *C. baccatum* var*. pendulum* UENF 1556 (Potnis *et al*. 2011). Among them, only *Bs2* and *Bs3*, and to some extent *Bs1*, have been commercially deployed. Based on gene-for-gene interactions between R genes and their corresponding avirulence genes, BSP causing *X. euvesicatoria* has been classified into eleven races (P0 – P10) (Stall *et al*. 2009). *Bs1* provides resistance against races P0, P2, and P5; *Bs2* against races P0, P1, P2, P3, P7, and P8; and *Bs3* against races P0, P1, P4, P7, and P9. Dominant resistance following infection often results in elicitation of a hypersensitive reaction (HR) and programmed cell death which creates high selection pressure for emergence and enrichment of pathogen races that overcome such resistance through loss/modification of avirulence genes (Gassmann *et al*. 2000). As a result, R genes are usually short-lived as exemplified by emergence and increased prevalence of races P6 and P10 in bell pepper cultivation, which are insensitive to the deployed R-genes (Kousik and Ritchie 1996a, b, 1998; Stall et al. 2009).

In contrast to R genes, recessive resistances typically result from the loss or modification of host susceptibility (S) factors that are exploited by bacteria to initiate a disease response (Sharma et al. 2022a). Recessive resistances are not race-specific and, following infection, do not elicit an HR — the lower selection pressure reduces the chance of emergence of virulent strains (Parlevliet 2002; Poland et al. 2009). This makes recessive resistance, despite the breeding challenges, highly desirable for management of rapidly evolving bacterial pathogens, such as *X. euvesicatoria*. Currently, three recessive resistances have been identified against BSP — *bs5* derived from *C. annuum* PI 271322, *bs6* from *C. annuum* PI 163192 or PI 264281, and *bs8* from *C. annuum* PI 163192 (Jones et al. 2002; Sharma et al. 2022b). Two of these genes, *bs5* and *bs6*, confer resistance to all known *Xe* races, including race P6 and P10 (Jones *et al*. 2002; Vallejos *et al*. 2010). Although *bs8* has been demonstrated to suppress *Xg*, its effect on *Xe* is not known (Sharma et al. 2022b). Only *bs5* has been commercially deployed (McCarthy 2011, 2012), and there have been no reports of its suppression by *X. euvesicatoria*.

Both *bs5* and *bs6* were first reported as monogenic, recessive, non-HR resistances against *X. euvesicatoria* race P6 (Jones et al. 2002). Both resistance are derived from hot pepper accessions collected from India and maintained at the Southern Regional Plant Introduction Station (SRPIS), GA (Jacobsen et al. 1982). *bs5* was reported to originate from *C. annuum* PI 271322, which carries resistance against BSP (Sowell and Dempsey 1977). The most probable source of *bs6* is PI 163192, which was used by Dempsey et al. (1981) to develop the C44 series, including the Pep13 line used by Jones et al. (2002) (Lane et al. 1997). *bs5* was transferred to the bell pepper *C. annuum* Early CalWonder (ECW) background by repeated backcrosses to generate ECW-50R (*syn*. ECW-44) line (Jones et al. 2002). Similarly, a NIL of ECW containing *bs6* has been named ECW-60R.

In order to understand the mechanism of resistance, it is often necessary to identify the underlying resistance gene. This is accomplished by gene mapping, which is the process of determining the physical location of a gene in the genome. Mapping of a resistance gene locus also enables use of linked molecular markers (in addition to phenotypic selection) to accelerate the breeding process by marker-assisted selection. Genotyping-by-sequencing (GBS) is a robust method of gene mapping which utilizes sequencing technology to discover molecular markers (such as SNPs and InDels) and genotypes the samples with those markers in a single step (Elshire et al. 2011). As a large number of small genomic variations from all chromosomes can be utilized in mapping, GBS is more precise and has higher resolving power than traditional genotyping methods. In this paper, we (i) identified the genomic localization of *bs5* and *bs6* resistance gene in pepper genome using GBS, (ii) fine mapped the respective resistance regions and identified flanking markers, and (iii) identified and analyzed candidate resistance genes.

## Materials and Methods

### Planting materials and growing conditions

For developing populations segregating for resistance, ECW50R and ECW60R were used as resistant parent for *bs5* and *bs6*, respectively. ECW was used as susceptible parent for both populations. For both resistances, ECW was crossed with respective resistant parent to produce an F_1_ population, which was self-pollinated to generate F_2_ seeds. F_3_ populations were generated by selfing of F_2_s when necessary. F_2_ recombinant individuals were self-pollinated, and progeny were genotyped to identify plants fixed for the recombined chromosomal segments (recombinant inbred lines (RILs)). A complete outline of all populations is presented in Figure S1. For all plants, seeds were sown in a seedling flat, and fourteen-day-old seedlings were transplanted to 10-cm pots containing Fafard Mix 4 (Fafard, Inc., Agawam, MA). For fine-mapping F2 populations, the plants were grown in 242-well trays (Speedling Inc., Sun City, FL) containing Speedling peat-lite soilless media (Speedling Inc., Sun City, FL). The transplants were grown in a greenhouse at temperatures ranging between 20-30°C.

### Inoculation and disease evaluation

As the resistant responses due to *bs5* and *bs6* does not result in HR induction, they are differentiated from the susceptible response by infiltration of bacterial suspension into pepper leaves at a low concentration (Stall 1981). In contrast to the development of necrotic lesion in susceptible pepper, the *bs5* resistance only causes a slight yellowing of the infiltrated area and the *bs6* resistant response is characterized by a more intense chlorosis (Vallejos et al. 2010). *X. euvesicatoria* race P6 strain Xv157 was grown in nutrient broth (BBL, Cockeysville, MD) overnight at 28°C with constant shaking. Bacterial cells were pelleted by centrifugation, the supernatant was discarded, and the cells were re-suspended in sterile tap water. The bacterial suspension was adjusted using Spectronic 20 Genesys spectrophotometer (Spectronic Instruments, Rochester, NY) to OD_600_=0.3, which is approximately 10^8^ CFU/ml, then diluted to 10^5^ CFU/ml in sterile tap water. The resulting bacterial suspension was infiltrated with a syringe and hypodermic needle into the mesophyll of the first and second true leaf of four-week-old pepper plants. Inoculated plants were maintained in a greenhouse for disease development, and the plants were evaluated three weeks after inoculation. Plants showing confluent necrosis were rated as susceptible, else they were rated as resistant for the respective resistance. For *bs6* resistance, the disease screen of each RIL was repeated multiple times to obtain accurate phenotypic result.

### GBS library preparation and sequencing

Foliar tissue from young leaves was lyophilized and used for DNA extraction. Genomic DNA was extracted using the Qiagen Plant DNeasy Mini Kit (Qiagen, Germantown, MD) according to the manufacturer’s instructions. The DNA was normalized to 5 ng/μL based on quantification with a Synergy 2 multimode microplate reader (Biotek Instruments, Winooski, VT) with the Quant-iT PicoGreen double-stranded DNA quantification assay (Thermo Fisher Scientific, Waltham, MA). A 96-plex (ninety one F_2_s, a single F_1_, and two each of ECW and respective resistant parent) *ApeKI* GBS library was constructed using a previously published protocol (Elshire et al. 2011). Barcode-adapter titration indicated that 0.9 ng μL^-1^ of each barcode-adapter per 50 ng of genomic DNA produced satisfactory libraries without dimer formation. The barcode-adapter titration mixture and the final GBS library were analyzed on an Agilent 2100 Bioanalyzer (Agilent Technologies, Santa Clara, CA) to ensure acceptable fragment size distribution and quantities. The GBS library was diluted to 3.6 pM and sequenced on one lane (single end, 101 base pair read length) of an Illumina HiSeq 2500 (Illumina Inc, San Diego, CA) at the Genomics Resources Core Facility (Weill Cornell Medicine, NY).

### GBS pipeline and SNP discovery

The raw sequencing reads were processed in TASSEL version 3.0 (Bradbury et al. 2007) using either the reference genome-reliant TASSEL-GBS pipeline (Glaubitz et al. 2014) or the reference-free UNEAK pipeline (for *bs6*) (Lu et al. 2013). For both pipelines, high quality sequencing reads that contained a barcode-adapter, an *ApeKI* restriction site, and an inserted genomic sequence (hereafter termed GBS tags) were identified and selected based on polymorphism between parents. In TASSEL-GBS pipeline, the reads were aligned with the Burrows-Wheeler Alignment (Li and Durbin 2009) to the *C. annuum* UCD10X reference genome, release 1.1 (Hulse-Kemp et al. 2018) to identify polymorphisms (Table S1). For the UNEAK pipeline, reference genome information was not necessary, and SNPs were identified by pairwise alignment of all unique sequence tags across the entire dataset (Table S2). Raw read files from sequencing of GBS libraries are deposited in NCBI SRA under bioproject <u>PRJNA863731</u>.

### Linkage analysis

Polymorphic SNPs identified between the parental lines were employed for linkage analyses using MapDisto v1.7 (implemented within Microsoft Excel 2007), (Lorieux 2012). The parameters in linkage analyses were a minimum LOD=5, a maximum r=0.3, and the ‘Kosambi’ mapping function. The loci were ordered within each linkage map using the auto-order function. QTL analysis was conducted for each population to determine the association between the SNPs within a linkage group and resistance to race P6. Single marker analysis was performed using the R/qtl package in R v3.3.1 (Broman et al. 2003).

### CAPS marker development and genotyping

Cleaved Amplified Polymorphic Sequence (CAPS) markers were designed for validating the mapping results from GBS and for fine mapping. Primers for the markers were designed using Primer 3 software (Untergasser et al. 2007) utilizing SNPs identified from GBS. DNA was extracted using a Cetyltrimethylammonium Bromide (CTAB) method (Doyle and Doyle 1987) and polymerase chain reaction (PCR) was carried out with Phire Hot Start II DNA polymerase (Thermo Fisher Scientific, Waltham, MA) in a 10 μl volume, which consisted of 2 μl of DNA (adjusted to ~20 ng/μl), 4.89 μl of HPLC-H_2_O, 2 μl of 5X Phire Reaction Buffer, 1 μl of dNTPs, 0.03 μl each of forward and reverse primers, and 0.05 μl of polymerase. The amplicons were digested with appropriate restriction enzymes according to the manufacturer’s recommendations (New England Biolabs, Ipswich, MA). Results were detected using electrophoresis on 3% agarose gels stained with ethidium bromide.

### HRM marker development and genotyping

High Resolution Melting curve (HRM) markers were developed from SNPs between purines and pyrimidines identified from GBS. Primers were developed using the IDT PrimerQuest (<u>idtdna.com/Primerquest</u>). DNA was extracted using a NaOH rapid DNA extraction method (Lee et al. 2017). The 5 μl PCR reactions were mixed with 2x AccuStart II PCR SuperMix (Quantabio, Beverly, MA), 0.5 μM of each primer, and 20x EvaGreen Dye (Biotium, Hayward, CA) and run as follows: (95°C @ 60s) + 40 × ((94°C @ 5s) + (*Tm* @ 10s) + (72°C @ 15s)) + (72°C for 60s), where *Tm* is the annealing temperature. For allele determination, melting curve analysis was performed by scanning the PCR product in a LightCycler 480 Instrument II (Roche, Pleasanton, CA).

### Whole genome sequencing

A modified microprep protocol was used for DNA extraction for whole genome sequencing (Fulton et al. 1995; Sharma et al. 2022b). DNA concentration and purity was verified using NanoDrop (Thermo Fisher Scientific, Waltham, MA). Subsequently, DNA was cleaned using DNeasy PowerClean Pro Cleanup Kit (Qiagen, Germantown, MD) following the manufacturer’s recommendations. Illumina sequencing library was prepared using a Nextera DNA Flex Library Prep Kit (Illumina Inc, San Diego, CA) using the protocol recommended by the manufacturer. The DNA was sequenced to produce 100 base-pairs (bp) paired end reads in one lane of Illumina HiSeq 3000 at University of Florida Interdisciplinary Center for Biotechnology Research.

### Super-scaffolding

The *C. annuum* ECW whole genome sequence (<u>GCA_011745845.1</u>) is only assembled to scaffold level at the time of analysis (Kim et al. 2017). To produce contiguous sequence, the *bs5* or *bs6* fine mapped intervals were blasted against the reference genome *C. annuum* UCD10X (<u>GCF_0028783951</u>). All ECW scaffolds with query coverage greater than 2% and matching to unique regions were identified and concatenated together in correct order and orientation to produce ECW super-scaffolds for *bs5* and *bs6*. The super-scaffolds also consisted of 5 Kbp region up- and down-stream from flanking markers and 3 Kbp gap between stitched scaffolds. The super-scaffolds were aligned with *C. annuum* UCD10X resistance intervals to verify complete coverage.

### Super-scaffold gene prediction

The ECW genes were predicted *de-novo* to overcome differences in gene annotations between reference genomes. ECW gene prediction model was developed using BRAKER v2.1.6 (Brůna et al. 2021). Within BRAKER, three publicly available ECW RNAseq sequences (<u>SRR13488414, SRR13488423</u>, and <u>SRR13488424</u>) were aligned to *C. annuum* ECW genome sequence (*GCA_011745845.1*) and supplied to Genemark-ET v4.68 (Lomsadze et al. 2014) to generate hints for training Augustus v3.4.0 (Stanke et al. 2008). The resulting ECW gene prediction model was used to identify potential protein coding regions in the *bs5* and *bs6* super-scaffolds. The genes were validated based on their posterior probability and annotation of homologous regions in *C. annuum* UCD10X or *C. annuum* CM334 annotation.

### Sequence analysis

Polymorphisms for *bs5* were identified using whole genome bulk sequences of PI 163192 × ECW50R F_2_ population, which is fixed for *bs5* gene (Sharma et al. 2022b). For *bs6*, the whole genome sequence of ECW60R was used. The sequences were analyzed using an in-house pipeline. The quality of the reads was verified with FastQC 0.11.7 (bioinformatics.babraham.ac.uk/projects/fastqc) and the adapters were trimmed using trim_galore v0.6.5 (Krueger et al. 2021). The trimmed reads were aligned to *C. annuum* ECW genome using Bwa-mem2 v2.2.1 (Vasimuddin et al. 2019). The resulting alignment file was used for variant calling with the HaplotypeCaller tool in GATK 4 (DePristo *et al*. 2011). The variants were filtered under high stringency as follows: depth ≥ 12, quality-normalized depth ≥ 10, mapping quality ≥ 50, and reference allele depth ≤ 0.1 * alternate allele depth. The sequencing data for PI 163192 × ECW50R F_2_s has previously been deposited in NCBI/ENA/DDBJ database under bioproject <u>PRJNA789991</u>. ECW60R whole genome sequence is deposited under bioproject <u>PRJNA863893</u>.

### Candidate genes identification

The coordinates and allelic sequence of high-quality polymorphisms in *bs5/bs6* super-scaffolds were derived from variant calling of *C. annuum* ECW scaffolds with an in-house script. The polymorphism were annotated with snpEff v5.0 (Cingolani *et al*. 2012) using a custom superscaffold variant annotation database built using previously described sequences and protein coding regions. Only the variations that result in protein coding changes were selected to identify potential candidate genes. Potential homologs of candidate genes in other *C. annuum* genomes were identified by blasting the predicted amino acid sequences of those genes, which also provided the functional annotations of the candidates. Finally, protein domains containing the polymorphisms between ECW and ECW50R/ECW60R were identified by Pfam (Mistry et al. 2021) and InterPro search (Blum et al. 2021).

## Results

### Segregation and phenotype

The phenotypic differences between ECW and ECW50R (*bs5*) were clear and easily distinguishable following inoculation at a relatively low bacterial concentration (10^5^ CFU/ml) (Figure 1). The ECW leaf tissue developed necrotic lesions surrounded by yellow halo while the ECW50R tissue remained mostly green. In the GBS F_2_ population, 91 out of 100 F_2_s (19 resistant and 72 susceptible) were phenotyped with high confidence and thus were used for GBS step. The ratio of resistant to susceptible F_2_s (1:3.8) was slightly lower than the expected ratio of 1:3 for recessive monogenic inheritance, however the difference was not statistically significant (*χ^2^=0.824* at 1 degree of freedom; *p=0.364*).

**Figure 1.**
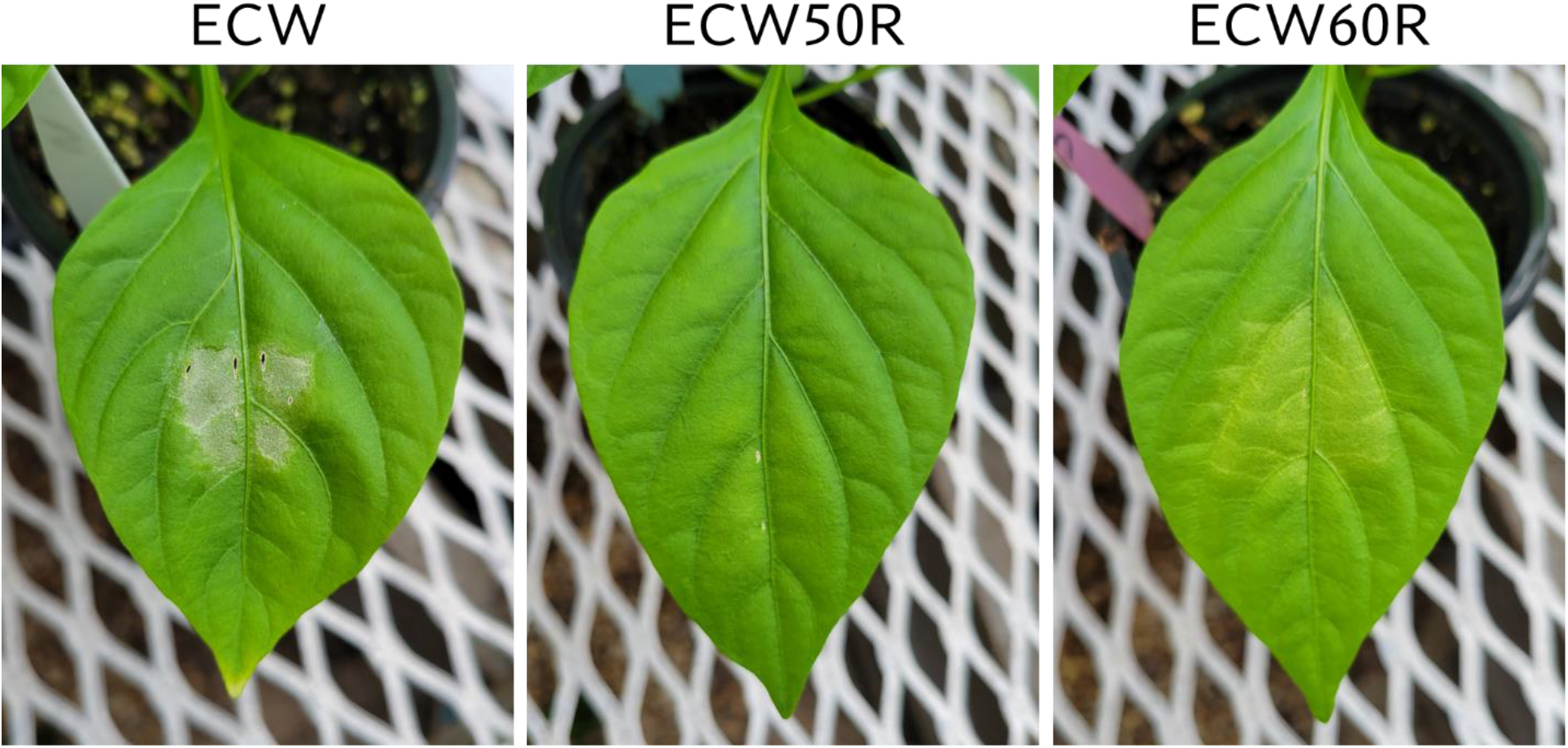
Phenotypes of ECW, ECW50R (*bs5*), and ECW60R (*bs6*) pepper 5 days after inoculation of *Xanthomonas euvesicatoria* strain Xv157 at 10^5^ CFU/ml.

The phenotype of ECW60R (*bs6*) resistance was not as distinct as *bs5* (Figure 1). As expected, *bs6* resistance was characterized by extensive chlorosis. Out of 120 F_2_s, 92 most clearly phenotyped individuals (29 resistant and 63 susceptible) were selected for GBS analysis. The ratio of resistant to susceptible F_2_s (1:2.2) was not statistically different (*χ^2^=2.087* at 1 degree of freedom; *p=0.1486*) from the expected 1:3 ratio.

### *bs5* locus is linked to shorter arm of chromosome 3

A total of 169,398,995 reads were generated from the *bs5* GBS library (Table S1). The GBS pipeline discovered 101 high quality SNPs that were polymorphic between the two parents, and those SNPs were selected for further analysis. The linkage analysis of 88 F_2_s that could be genotyped identified thirteen linkage groups, and the *bs5* resistance mapped to linkage group 1 in chromosome 3 with highest significance (Figure 2; Table S3; Table S4). SNPs between positions 134,620 and 1,098,542 of chromosome 3 were the most significantly associated with *bs5* (p<0.0001). Genotyping of the F2 population with CAPS markers spanning the linkage region confirmed 100% marker-trait co-segregation in the mapping population (Table 1; Table S5). The results indicate that *bs5* is located towards the distal end of the short arm of chromosome 3, within a ~1 Mbp interval between 0.1 and 1.1 Mbp position.

**Figure 2.**
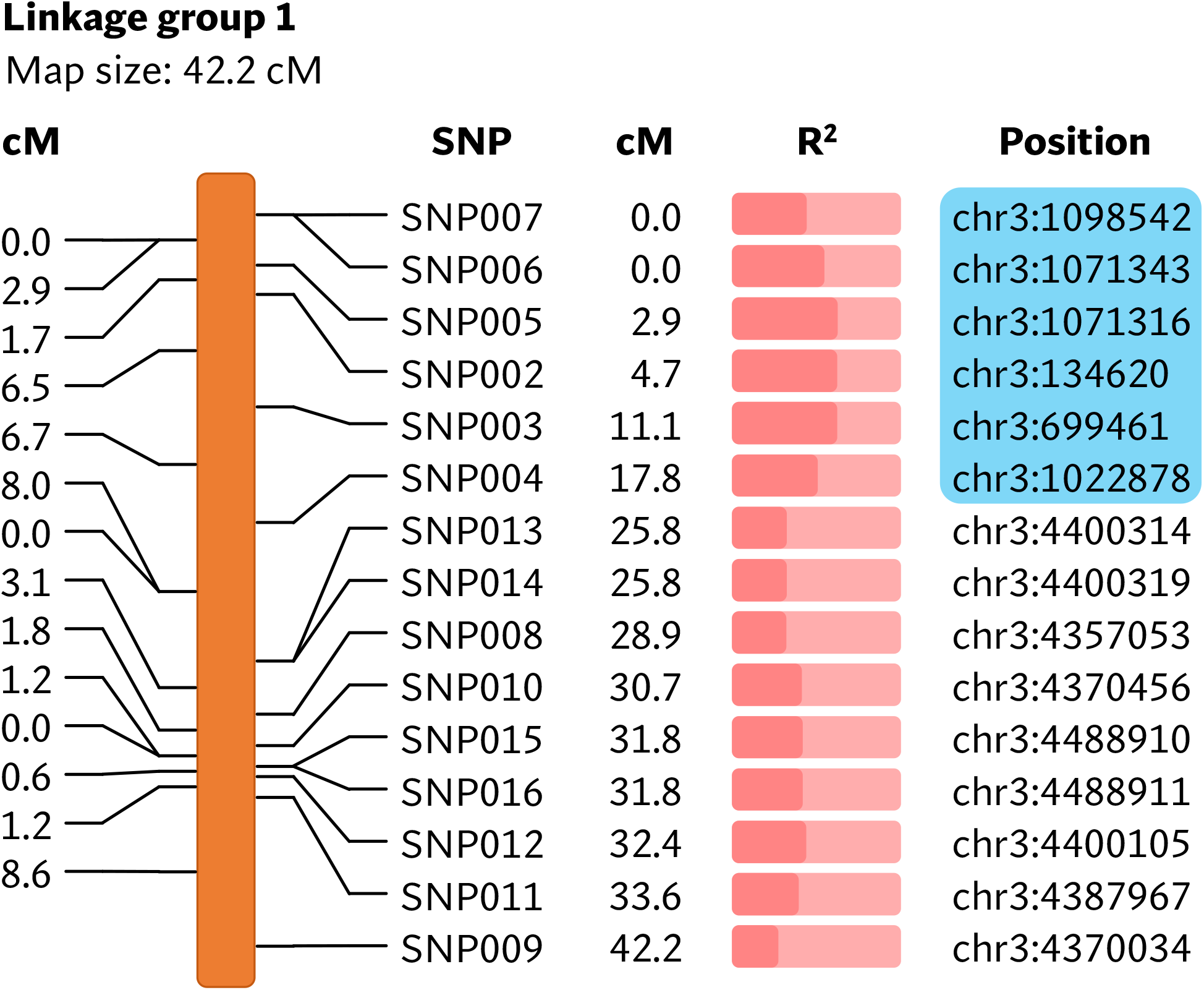
Linkage map showing markers associated with *bs5*. The physical positions of markers are based on *C. annuum* UCD10X genome, release 1.1. Blue box encloses genomic area that was further investigated by fine-mapping.

**Table 1.**
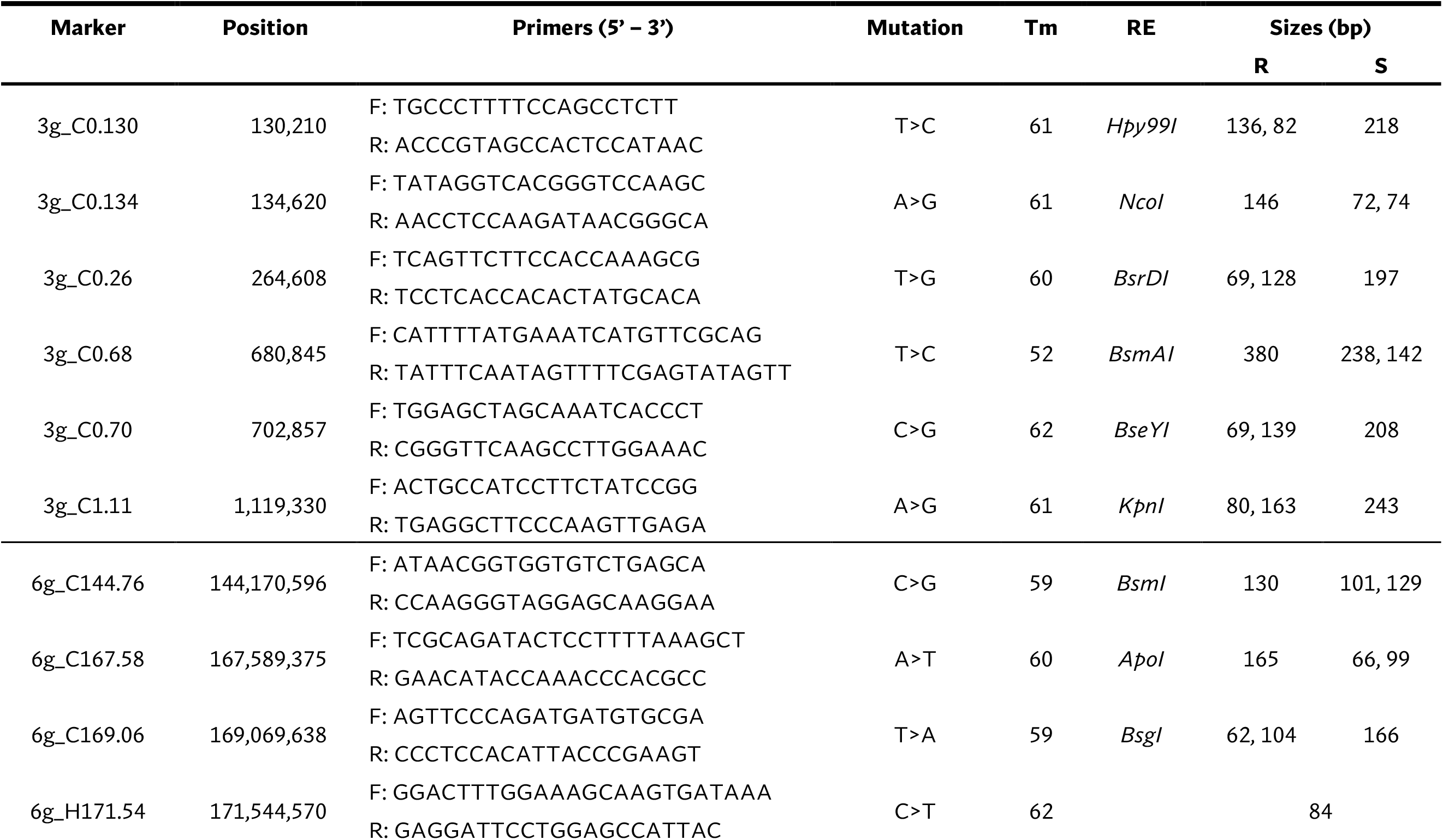
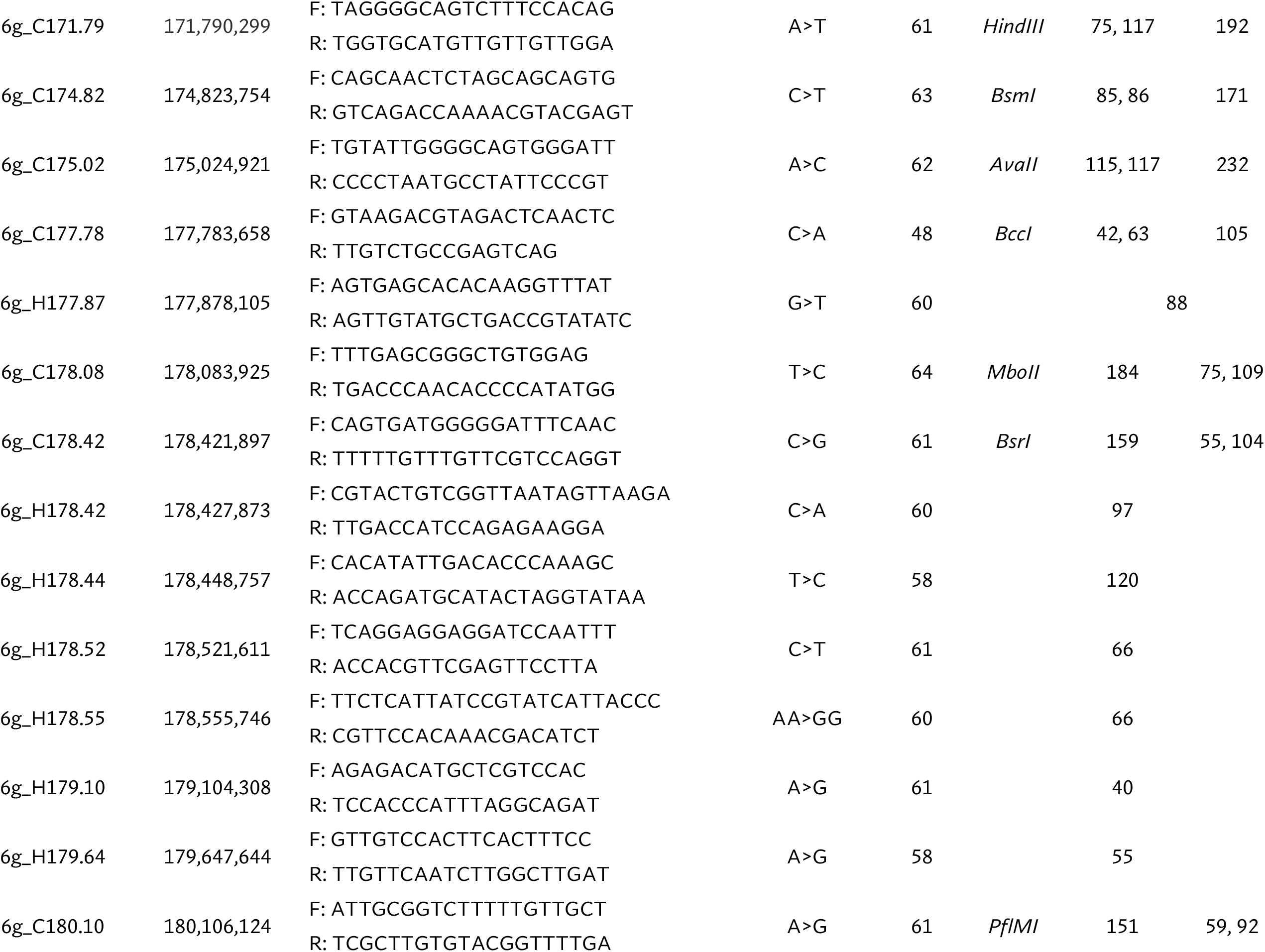
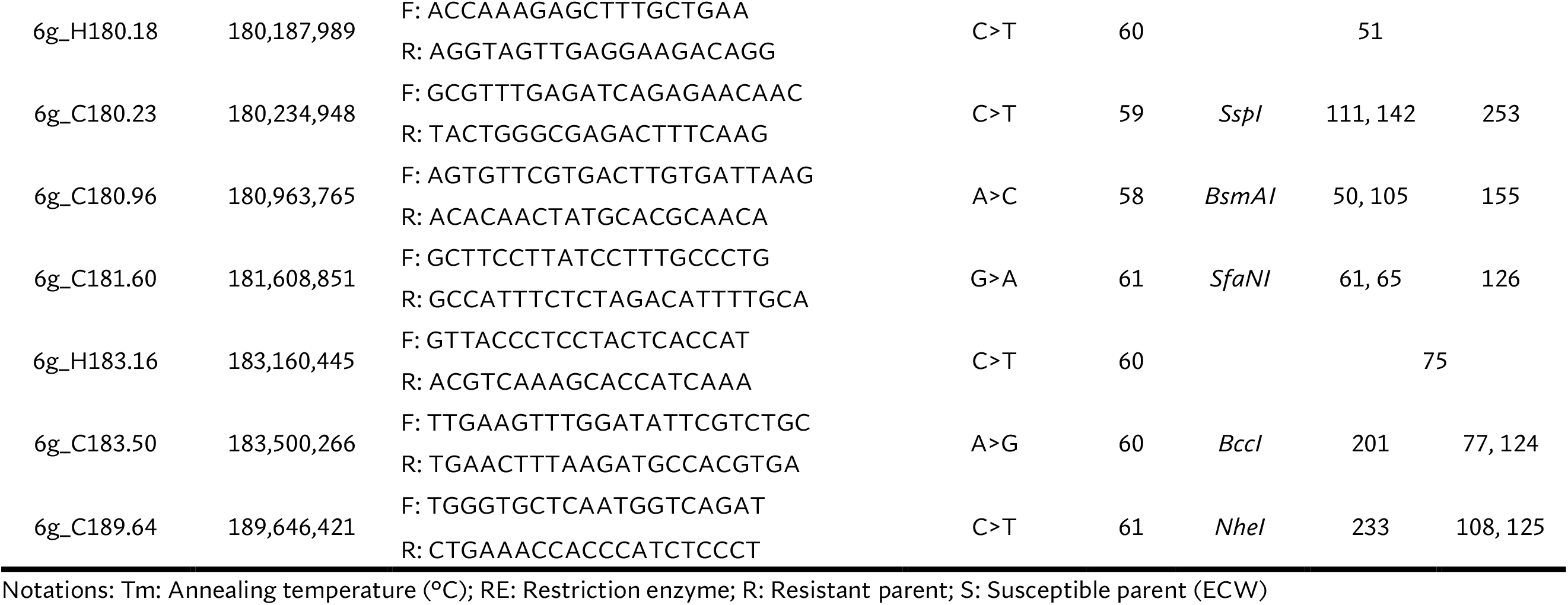
Molecular markers used for fine mapping the position of *bs5* and *bs6* resistance. Markers starting with ‘3g’ are located in chromosome 3 and linked to *bs5*, whereas those starting with ‘6g’ are located in chromosome 6 and linked to *bs6*. ‘C’ and ‘H’ in the name indicate that they are CAPS and HRM markers, respectively. Genomic positions are based on *C. annuum* UCD10X reference genome release 1.1.

### *bs5* is fine-mapped to a 546 Kbp interval in chromosome 3 telomere

A larger ECW × ECW50R F_2_ population was developed to fine-map the position of *bs5*. Out of 1270 F_2_s genotyped with flanking markers 3g_C0.134 and 3g_C1.11, 16 individuals were identified as recombinants and were phenotyped. Ten informative recombinants and F3 RILs developed from six non-informative recombinants placed *bs5* into an ~546 Kb interval between markers 3g_C0.134 (~0.4 cM) and 3g_C0.68 (~0.95 cM) with tight linkage with marker 3g_C0.26. (Figure 3; Table S6. Genotyping results for recombinant progenies from ECW × ECW50R fine-mapping F_2_ population. Table S7; Table S7).

**Figure 3.**
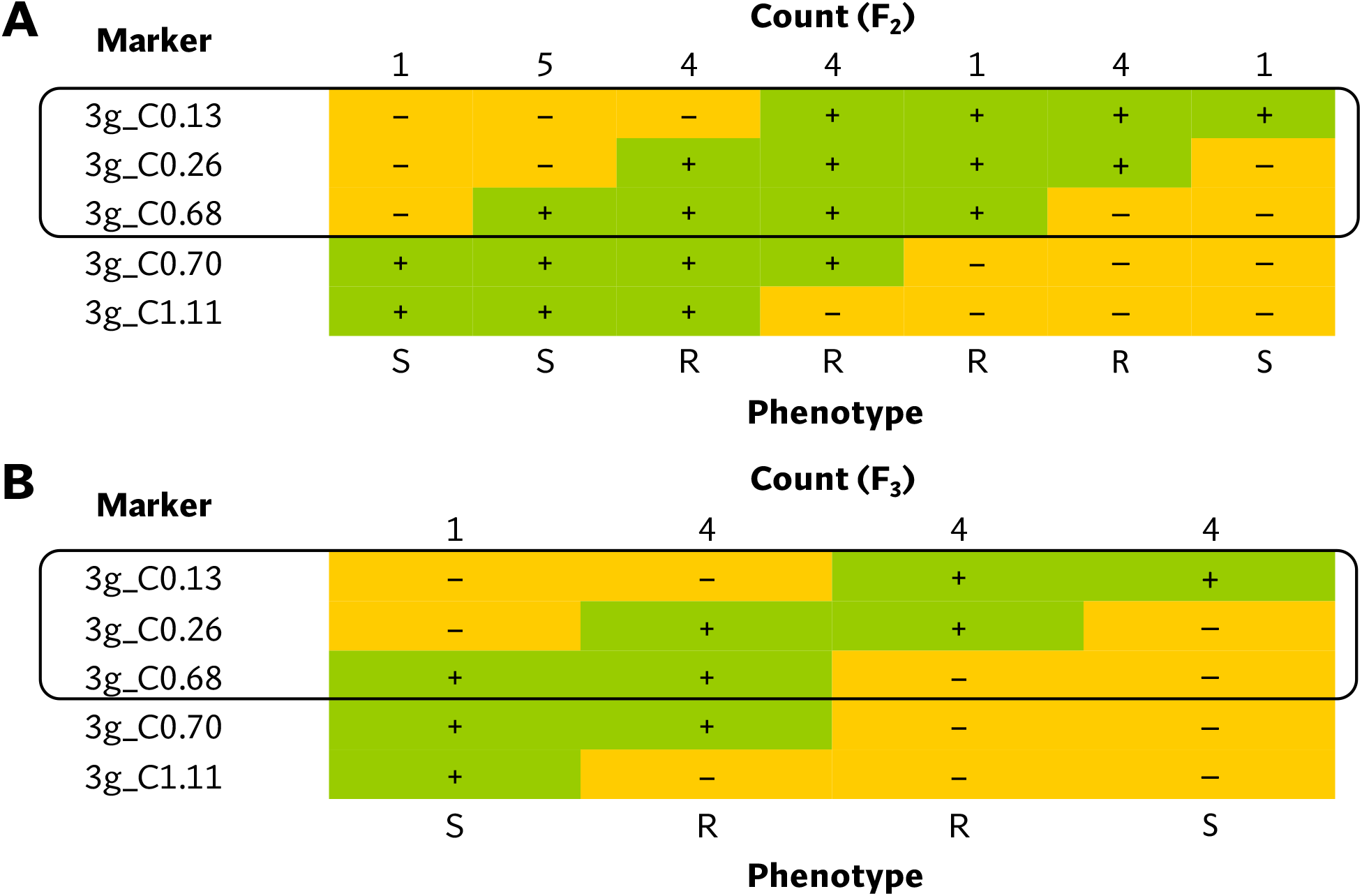
Tabulation of genotypes of the (A) F2 and (B) F3 progenies from *bs5* fine-mapping population that recombine within the *bs5* mapped region, together with their phenotypes. The black box encloses the closest markers flanking the new resistance interval. Notations: +: homozygous for the resistant/ECW50R allele; -: heterozygous or homozygous for the susceptible/ECW allele; R: resistant phenotype; S: susceptible phenotype

### *bs5* interval contains 15 polymorphic candidate genes

An ECW *bs5* super-scaffold was developed by concatenating *C. annuum* ECW scaffolds that align with in *C. annuum* UCD10X *bs5* interval. This super-scaffold consisted of 535 Kbp sequence including gaps and flanking region and provided complete coverage of UCD10X *bs5* interval (Table S8). Comparison of whole genome polymorphisms between *bs5*-fixed line (PI 163192 × ECW50R) and ECW identified a total of 1,718 variations in this region under stringent filtration (data not shown). However, only 31 variations were found to alter the proteins sequences, which resulted in 14 putative candidate genes for *bs5* resistance (Table 2).

**Table 2.**
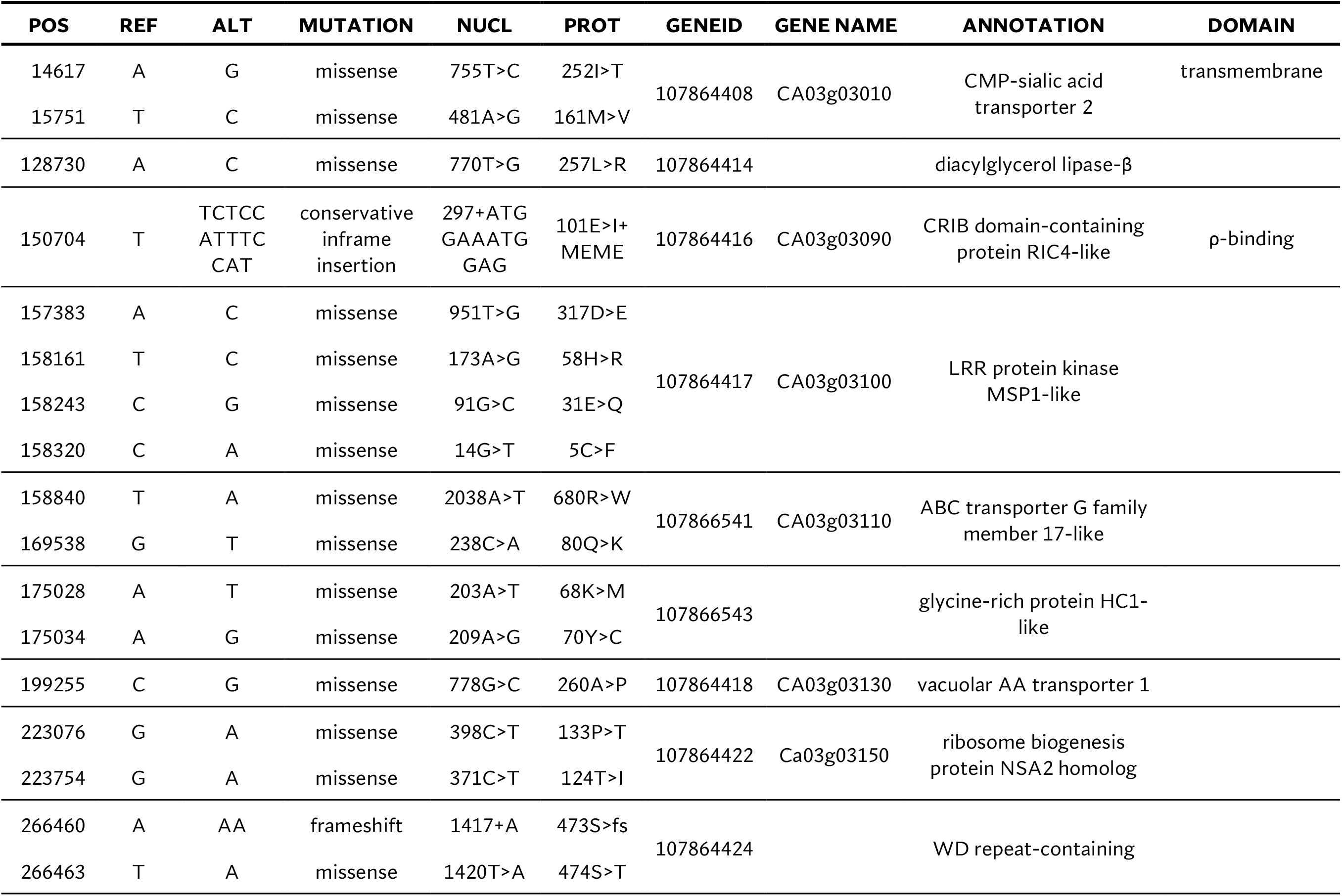
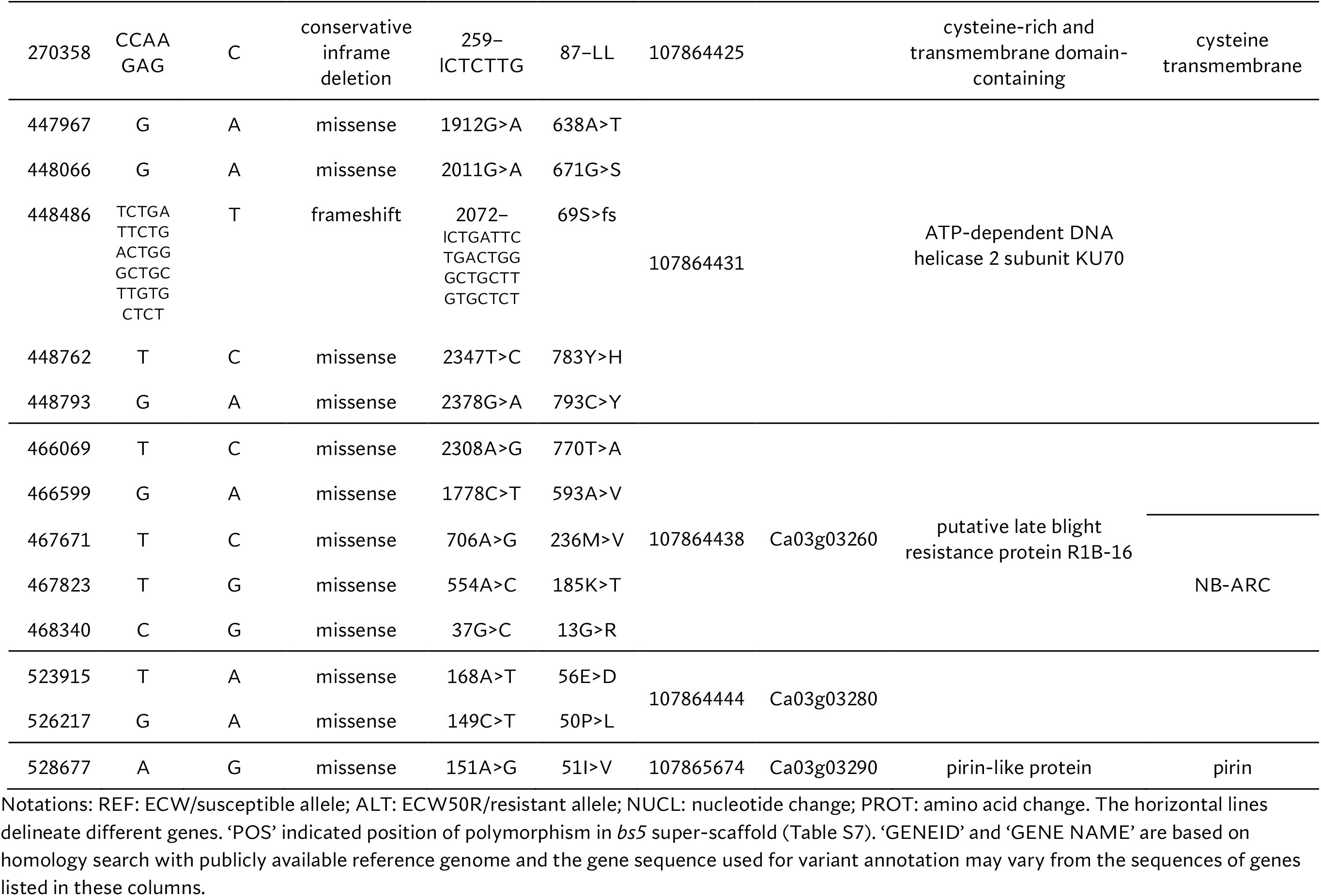
List of candidate genes for *bs5* resistance.

### *bs6* locus is located in chromosome 6

As the reference-based GBS pipeline only identified a small number of polymorphic markers, the reference-free UNEAK pipeline was used for mapping *bs6*. This pipeline discovered 133 SNPs from a total of 173,074,228 reads generated from sequencing (Table S2). Nine linkage groups were generated from the linkage analysis using genotyping information from 92 F_2_ plants (Table S9), out of which the *bs6* resistance phenotype was significantly (p < 0.0001) linked to SNPs on linkage group 3 (Figure 4; Table S9; Table S10). The linkage group was determined to be physically located in chromosome 6. CAPS makers were developed in the *bs6*-mapped region, and genotyping of the F_2_ population validated the linkage between those markers and the resistance phenotype (Table 1; Table S11). The results indicated that *bs6* was located within an ~21 Mbp interval between positions 168–189 Mbp in *C. annuum* UCD10X genome.

**Figure 4.**
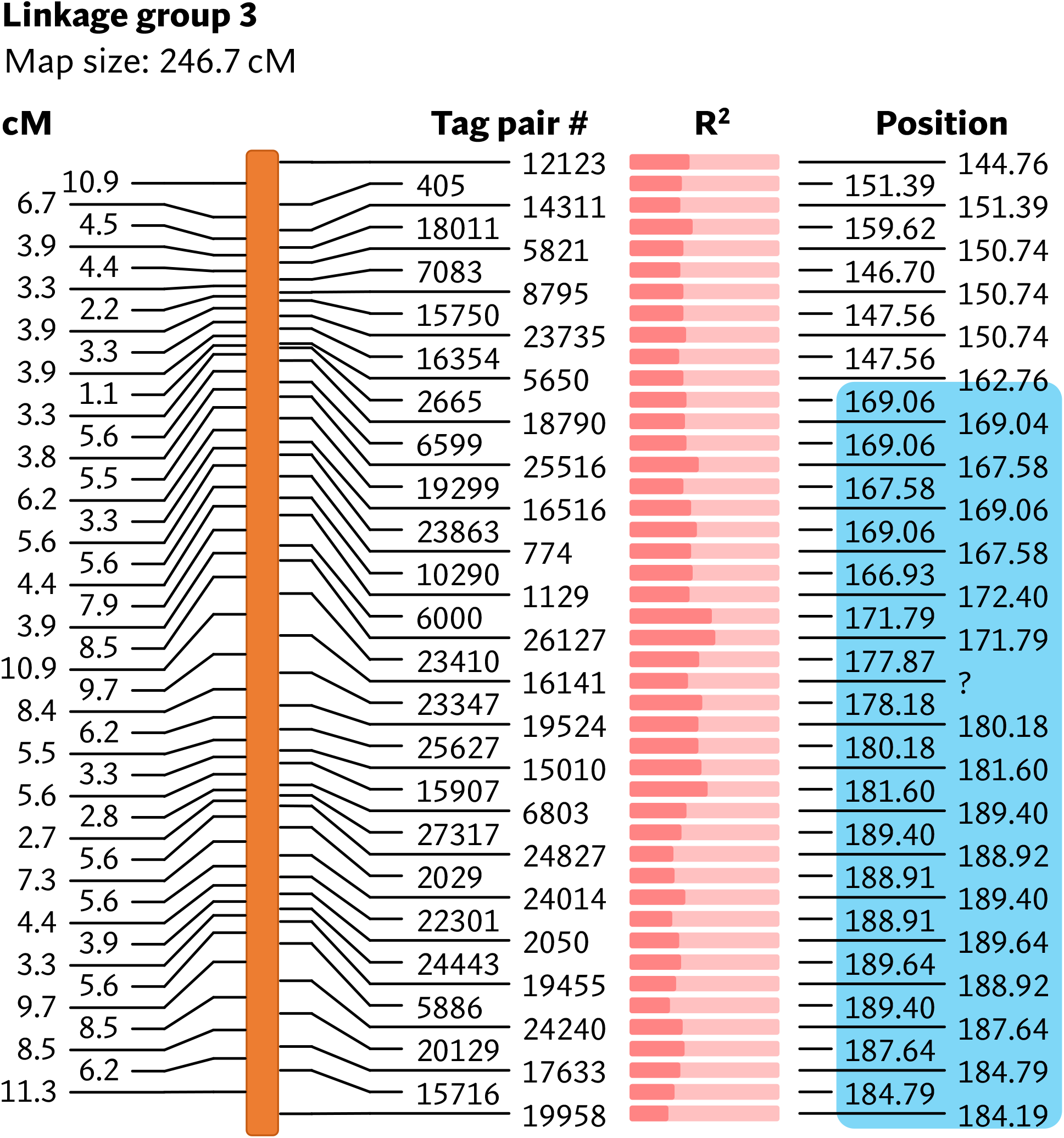
Linkage map showing markers associated with *bs6*. The physical positions of the markers are based on *C. annuum* UCD10X genome, release 1.1. Blue box encloses genomic area that was further investigated by fine-mapping.

### *bs6* is fine-mapped to a 656 Kb interval

Five of the CAPS markers within the ~21 Mbp *bs6* interval were initially used to more precisely determine the position of *bs6*. In a fine mapping F_2_ population of 940 plants, 277 plants were identified as recombinants, 123 of which were homozygous for 60R alleles throughout part of the recombined region and were phenotyped as F2 plants; genotyping of these F_2_s delimited the resistance locus to an ~9.8 Mbp region between markers 6g_C171.79 and 6g_C181.60 (Figure 5A; Table S12). F_3_ RILs developed from 61 F_2_s that recombined within the region were genotyped with eight new CAPS markers within the interval (Table 1); this delimited *bs6* within an ~5.1 Mbp interval between markers 6g_C175.02 and 6g_C180.10 (Figure 5B; Table S13). A second ECW60R × ECW F_2_ population of 940 plants was developed and genotyped with new HRM markers (Table 1), and 41 recombinants between flanking markers 6g_H171.54 and 6g_H183.16 were identified and developed into F3 RILs. All 41 RILs were phenotyped and were genotyped with markers in the 5.1 Kbp interval, thereby delimiting *bs6* to an ~656 Kbp region between markers 6g_H178.44 (~0.11 cM) and 6g_H179.10 (~0.11 cM) (Figure 5C; Table S14).

**Figure 5.**
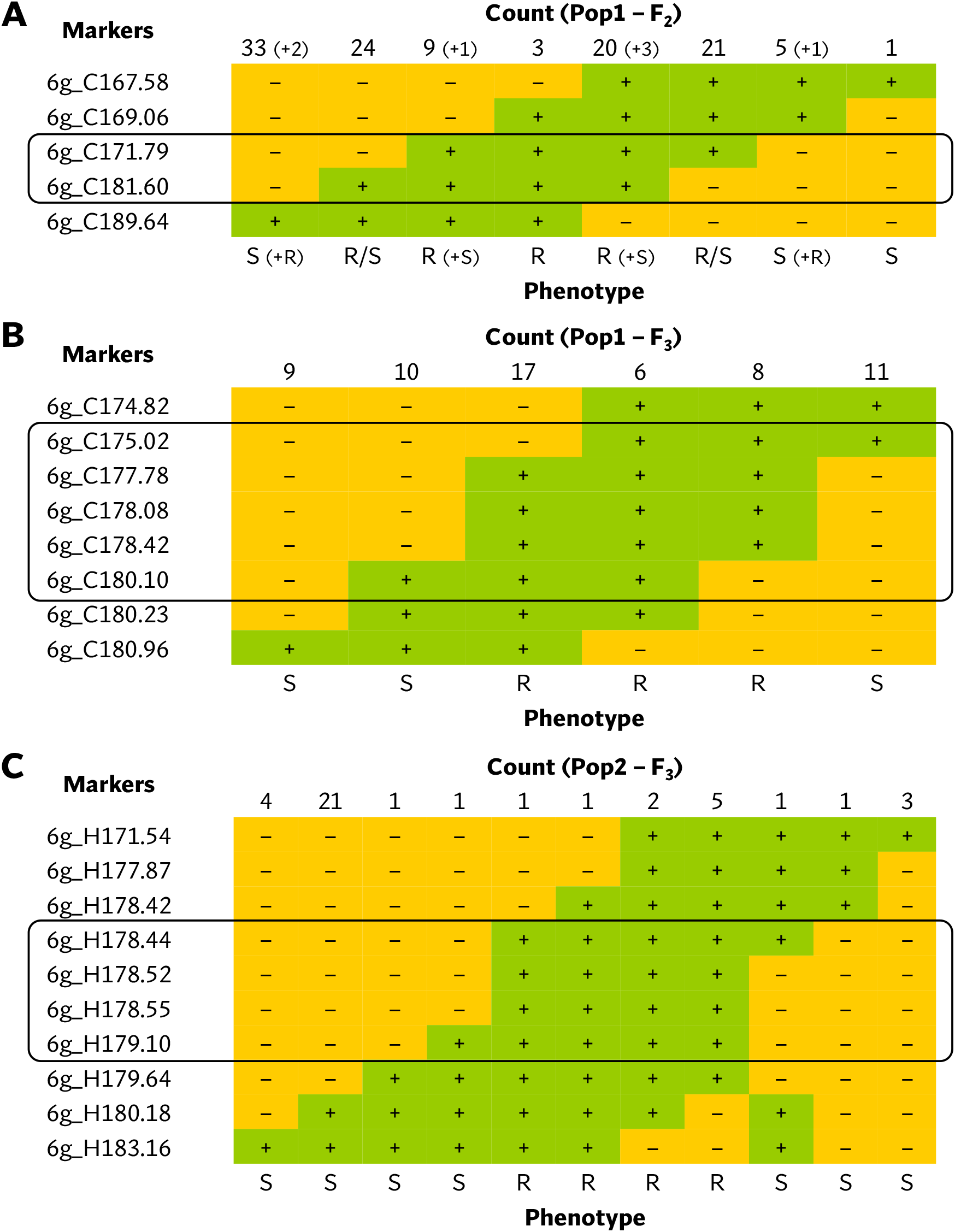
Tabulation of genotypes of the (A) F2 plants from first *bs6* fine-mapping population, (B) F3 plants from first *bs6* fine-mapping population, and (C) F3 plants from second *bs6* fine-mapping population that recombine within the *bs6* interval, together with their phenotypes. The black boxes enclose the closest markers flanking the new resistance interval. Notations: +: homozygous for the resistant/ECW60R allele; -: heterozygous or homozygous for the susceptible/ECW allele; R: resistant phenotype; S: susceptible phenotype.

### *bs6* interval contains 8 polymorphic candidate genes

The ECW *bs6* super-scaffold spanned three *C. annuum* ECW scaffolds with a total size of 681 Kb, providing complete coverage of UCD10X *bs6* interval (Table S6. Genotyping results for recombinant progenies from ECW × ECW50R fine-mapping F_2_ population.

Table S7. Genotyping results for recombinant progenies from ECW × ECW50R fine-mapping F_3_ population. Table S8). A total of 1,718 variations were identified between ECW and ECW60R genome in this region after filtration. Annotation of those variations identified protein coding changes in eight genes, which are candidates for *bs6* (Table 3). Interestingly, four of those candidates are functionally annotated as ZED1-related serine/ threonine kinases, and three have protein polymorphisms within the putative kinase domain (Table 3).

**Table 3.** List of candidate genes for *bs6* resistance.

| POS | REF | ALT | MUTATION | NUCL | PROT | GENEID | GENE NAME | ANNOTATION | DOMAIN |
| --- | --- | --- | --- | --- | --- | --- | --- | --- | --- |
| 2955 | T | A | missense | 887A>T | 296E>V | 107874896 | Ca06g12480 | Formyltetrahydrofolate<br>deformylase | Coiled coil |
| 4324 | G | A | stop gained | 439C>T | 147Q>* |  |  |  |  |
| 24554 | G | A | missense | 211C>T | 71L>F | 107872943 | FXO38_32047 | Phosphatidylserine<br>decarboxylase proenzyme 1 |  |
| 49875 | C | A | missense | 255C>A | 85D>E | 107872942 | Ca06g12500 | ZED1-related kinase (ZRK) 4 | S/T Kinase |
| 49930 | A | AT | frameshift | 311+T | 105A>fs |  |  |  |  |
| 50091 | T | A | missense | 471T>A | 157N>K |  |  |  |  |
| 52304 | T | TA | frameshift | 111+A | 38E>fs | 107874893 | Ca06g12510/<br>Ca06g12520 | ZRK1-like serine/ threonine-<br>protein kinase | S/T Kinase |
| 52358 | C | T | missense | 164C>T | 55S>F |  |  |  |  |
| 52928 | T | G | missense | 457T>G | 153S>A |  |  |  |  |
| 59302 | C | T | missense | 168G>A | 56M>I |  | FXO38_32052 | TCP-1/Cpn-60 chaperonin-like | Cpn-60 |
| 59330 | G | A | missense | 140C>T | 47S>F |  |  |  |  |
| 92420 | A | T | missense | 782A>T | 261E>V | 107874060 | Ca06g12530 | ZRK1-like serine/ threonine-<br>protein kinase |  |
| 92548 | C | G | missense | 910C>G | 304P>A |  |  |  |  |
| 92596 | C | A | missense | 958C>A | 320P>T |  |  |  |  |
| 94020 | G | A | missense | 122G>A | 41G>D |  | Ca06g12540 | Ubiquitin conjugating enzyme<br>variant (UEV) 1C-like | UBC_E2 |
| 94033 | C | G | missense | 135C>G | 45I>M |  |  |  |  |
| 94050 | C | G | missense | 152C>G | 51T>S |  |  |  |  |
| 94053 | A | G | missense | 155A>G | 52D>G |  |  |  |  |
| 127705 | TTA<br>A | T | disruptive<br>inframe<br>deletion | 308-ATA | 103-N | 107874895 | Ca06g12550 | ZED1-related kinase (ZRK) 1 | S/T Kinase |

## Discussion

In this paper, we show the genomic localization of two recessive BSP resistance genes: *bs5* and *bs6*. *bs5* was mapped to the telomeric region of chromosome 3 and *bs6* to chromosome 6. The genomic position of *bs5* is in discordance with a previous report on the position of *bs5*, which had mapped it to the centromeric region of chromosome 6 (Vallejos et al. 2010). However, the former positions were based upon two populations of 60 F_2_ and 88 F_3_ progenies and only utilized 64 markers for screening the entire pepper genome. In contrast, the *bs5* and *bs6* locations identified in the present study are based on mapping in F_2_ populations, were validated in large fine mapping populations, and benefited from larger numbers of markers identified through GBS. Furthermore, the availability of a high-quality pepper reference genome enabled us to cross-validate our GBS results.

Several pepper lines have been reported to have varying degrees of recessive resistance against BSP. One of the earliest discoveries recessive resistance was made by Dempsey (1953) in the pepper cultivar, Santanka. Hibberd *et al*. (1988) reported quantitative non-race-specific resistance in PI 163189. Poulos *et al*. (1992) reported that the quantitative, non-HR, non-race-specific resistance in CNPH 703 is controlled by at least two genes. Both PI 163189 and PI 183441 (parent of CNPH 703) were imported together with PI 163192, and thus the resistances in those accessions could also be due to *bs5/bs6*. A monogenic, recessive, non-HR and non-race-specific resistance in PI 163192 was identified by Szarka and Csilléry (2001) and named *gds* (general defense system); *gds* has since been shown to be the same as *bs5* (Timár et al. 2019). Riva et al. (2004) reported recessive resistance in UENF 1381 that may be governed by multiple genes. Furthermore, several genes have been identified in pepper which are required for complete virulence; reduced expression of such genes resulted in reduced susceptibility to BSP. Some notable examples include GLIP1 (Hong et al. 2008), MRP1 (An et al. 2008), MLO2 (Kim and Hwang 2012), and GRP1 (Kim et al. 2015).

A patent filed in 2013 and granted in the US in 2021 describes a recessive, non-race-specific resistance gene in pepper called “xcv-1”, which encodes a cysteine-rich transmembrane region with the resistant allele containing a double leucine deletion (Kiss et al. 2021). Interestingly, one of the polymorphic genes located towards the center of the *bs5* fine mapped interval (GeneID: 107864425) encodes a cysteine-rich transmembrane domain-containing protein (CYSTM) and has a double leucine deletion in the resistant allele (Table 2). The genomic localization of *xcv-1* has not been reported; however, out of 6 cysteine-rich transmembrane genes annotated in the *C. annuum* UCD10X genome, two are present in the *bs5* region (Table S15), and only 107864425 is polymorphic between ECW and ECW50R with a double leucine deletion (Table 2). Thus, it is likely that *xcv-1* and *bs5* are identical resistances (Szarka et al. 2022) and are encoded by gene 107864425. CYSTM proteins are known to have a role in stress tolerance and disease resistance. Ectopic overexpression of a group of pathogen-induced CYSTM proteins in *Arabidopsis* reduced in-planta population of *Pseudomonas syringae* pv. *tomato* (Pereira Mendes et al. 2021).

A number of *bs6* candidate resistance genes are ZED1-related kinases (ZRKs), which are members of the broad receptor-like kinase/Pelle family of protein kinases (Shiu et al. 2004). ZRKs belong to family RLCK-XII, which includes several pseudokinases that can participate in biotic defense response (Lewis et al. 2013; Wang et al. 2015; Seto et al. 2017). A tomato ZRK, *JIM2 (RxopJ4*), provides resistance against bacterial spot of tomato by serving as a decoy target for the type III effector, XopJ4, and consequently activates a ZAR1-mediated defense response (Schultink et al. 2019). Surprisingly, *RxopJ4* is one of several ZRKs located in the syntenic region of *bs6* in tomato genome (data not shown) (Sharlach 2013). Since ZRKs can be targeted by bacterial effectors, and since recessive resistances such as *bs6* often result from modification of bacterial susceptibility targets, four ZRKs in the *bs6* interval are also intriguing candidates for *bs6*.

*bs5* and *bs6* act synergistically and provide resistance against all races of *Xe*. Together with *bs8*, which provides resistance against *Xg*, they enable development of pepper varieties carrying long-lasting recessive resistance to all known BSP pathogens. Pyramiding of resistance genes also increases stability of resistance, both in terms of durability, and against unfavorable conditions. As an example, *bs5* or *bs6*, alone, provide lower of resistance at high temperatures (Vallejos et al. 2010). The next steps are to functionally characterize the candidate genes to identify *bs5/bs6*. Identification of the resistance genes will facilitate understanding of the mechanism of resistance, which in turn can contribute to the development of novel disease control strategies. Apart from pepper, development of bacterial spot-resistant tomatoes is highly desirable, and identification of the *bs5/bs6* genes will be a crucial step for identifying tomato homologs which can be targeted by gene-editing technologies.

## Acknowledgements

The research was supported by USDA NIFA Specialty Crop Research Initiative Grants Program Grant Number: 2015-51181-24312. We would also like to thank 2blades foundation for providing support for initial stages of mapping. *Xanthomonas euvesicatoria* strain Xv157 (race 6) was kindly provided by S. A. Miller (The Ohio State University, Wooster).

## Author contributions

G.V.M, J.B.J., and R.E.S. initially developed ECW50R and EW60R lines. J.P.H., and M.R.M. performed GBS sequencing and analysis. Fine mapping was conducted by J.L., R.W. and S.F.H. (genotyping) and G.V.M. and J.B.J. (phenotyping). U.S.G. generated the whole genome sequences. A.S. contributed to manuscript writing, fine-mapping, sequence analysis, and identification of candidate genes. All authors reviewed and revised the manuscript.

## Data availability

Raw read files from sequencing of GBS libraries are deposited in NCBI SRA under bioproject <u>PRJNA863731</u>. ECW50R F_2_s bulked sequence has previously been deposited in NCBI SRA under bioproject <u>PRJNA789991</u>. ECW60R whole genome sequence is deposited under bioproject <u>PRJNA863893</u>.

## Supplementary tables

(See supplementary excel file)

Table S1. Outline of TASSEL 3.0 Genotyping by Sequencing (GBS) pipeline for analysis of *bs5* raw data.

Table S2. Outline of TASSEL 3.0 Universal Network Enabled Analysis Kit (UNEAK) pipeline for analysis of *bs6* raw data.

Table S3. Genotypes of 88 (ECW x ECW50R) F2 plants at 121 SNPs identified by GBS, and phenotype of these plants upon infiltration with *X. euvesicatoria* Xv157.

Table S4. Single Marker Analysis of polymorphisms between ECW and ECW50R identified by Genotyping-by-Sequencing.

Table S5. Confirmation of *bs5* mapping in 88 ECW x ECW50R F2 plants with six CAPS markers located within the mapped region.

Table S6. Genotyping results for recombinant progenies from ECW × ECW50R fine-mapping F2 population.

Table S7. Genotyping results for recombinant progenies from ECW × ECW50R fine-mapping F3 population.

Table S8. Positions and orientations of *C. annuum* ECW scaffolds include in *bs5* (top) and *bs6* (bottom) super-scaffolds.

Table S9. Genotypes of 92 (ECW x 60R) F2 plants at 133 SNPs identified by GBS, and the phenotype of these plants upon infiltration with *X. euvesicatoria* race 6 (strain Xv157).

Table S10. Single Marker Analysis of polymorphisms between ECW and ECW60R identified by Genotyping-by-Sequencing.

Table S11. Confirmation of *bs6* mapping in 92 ECW x ECW60R F2 plants with six CAPS markers located within the mapped region.

Table S12. Genotyping results for recombinant progenies from the first ECW × ECW60R fine-mapping F2 population.

Table S13. Genotyping results for recombinant progenies from the first ECW × ECW60R fine-mapping F3 population.

Table S14. Genotyping results for recombinant progenies from the second ECW × ECW60R fine-mapping F3 population.

Table S15. List of cysteine-rice transmembrane proteins annotated in *C. annuum* UCD10X genome.

## Supplementary figures

**Figure S1.**
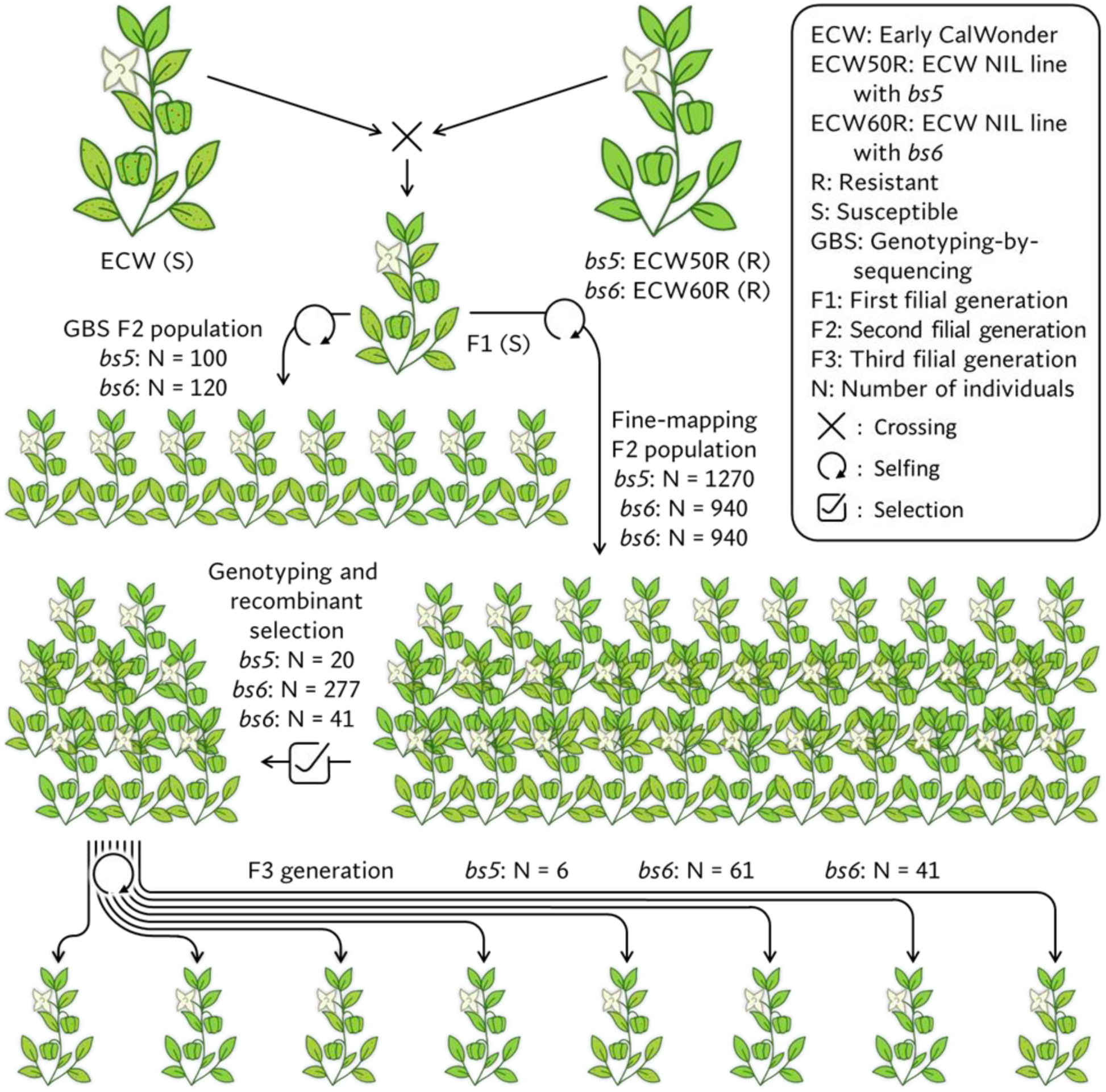
Schematic representations of crosses made to generate GBS and fine-mapping F2 population for *bs5* and *bs6*.

